# An AI-based segmentation and analysis pipeline for high-field MR monitoring of cerebral organoids

**DOI:** 10.1101/2023.04.07.535822

**Authors:** Luca Deininger, Sabine Jung-Klawitter, Petra Richter, Manuel Fischer, Kianush Karimian-Jazi, Michael O. Breckwoldt, Martin Bendszus, Sabine Heiland, Jens Kleesiek, Ralf Mikut, Daniel Hübschmann, Daniel Schwarz

## Abstract

**Background:** Cerebral organoids simulate the structure and function of the developing human brain *in vitro*, offering a large potential for personalized therapeutic strategies. The enormous growth of this research area over the past decade with its capability for clinical translation makes a non-invasive, automated analysis pipeline of organoids highly desirable.

**Purpose:** This work presents the first application of magnetic resonance imaging (MRI) for the non-invasive quantification and quality assessment of cerebral organoids using an automated analysis tool. Three specific objectives are addressed, namely organoid segmentation to investigate organoid development over time, global cysticity classification, and local cyst segmentation.

**Methods:** Nine wildtype cerebral organoids were imaged over nine weeks using high-field 9.4T MRI including a 3D T2*-w and 2D diffusion tensor imaging (DTI) sequence. This dataset was used to train a deep learning-based 3D U-Net for organoid and local cyst segmentation. For global cysticity classification, we developed a new metric, *compactness*, to separate low- and high-quality organoids.

**Results:** The 3D U-Net achieved a Dice score of 0.92±0.06 (mean ± SD) for organoid segmentation in the T2*-w sequence. For global cysticity classification, *compactness* separated low- and high-quality organoids with high accuracy (ROC AUC 0.98). DTI showed that low-quality organoids have a significantly higher diffusion than high-quality organoids (p < .001). For local cyst segmentation in T2*-w, the 3D U-Net achieved a Dice score of 0.63±0.15 (mean ± SD).

**Conclusion:** We present a novel non-invasive approach to monitor and analyze cerebral organoids over time using high-field MRI and state-of-the-art tools for automated image analysis, offering a comparative pipeline for personalized medicine. We show that organoid growth can be monitored reliably over time and low- and high-quality organoids can be separated with high accuracy. Local cyst segmentation is feasible but could be further improved in the future.

## 1. Introduction

Cerebral organoids are key models to study human brain tissue and probe pathophysiological processes with tremendous potential for tailored therapeutic strategies. They are patient-derived miniature 3D tissue cultures that are grown from induced pluripotent stem cells. Cerebral organoids have been used to study a wide range of neurological disorders like microcephaly [1] or neurodegenerative diseases like Alzheimer’s [2] or Parkinson’s disease [3].

The growing interest in organoid research over the past decade [4] results in an increasing amount of data and thus calls for automated analysis and quantification. However, current automated organoid analysis pipelines are limited to smaller, e.g. intestinal, organoids [5] or require organoid sacrifice [6]. Magnetic resonance imaging (MRI) allows for the generation of 3D cerebral organoid time series due to its non-invasive imaging procedure. Furthermore, brain MRI is the gold standard for diagnosis, staging, and treatment guidance of various neurological disorders, thus highlighting its potential for imaging cerebral organoids, which has not yet been exploited.

In the complex process of organoid cultivation, an important undesired route of organoid differentiation is marked by the occurrence of fluid-filled cavities (or ‘cysts’) [7, 8]. Thus, accurately and automatically estimating organoid cysticity would greatly contribute to organoid quality monitoring. So far, however, only an approach for automated segmentation of exophytic cysts in patients with polycystic kidney disease using MRI has been reported [9].

Here, we present the first application of MRI to human brain organoids using a neural network-based approach to extract cerebral organoid volume and structural features. Specifically, we address three crucial tasks for organoid monitoring and quality assessment: organoid segmentation, global cysticity classification, and local cyst segmentation.

## 2. Results

### Organoid segmentation

Organoid segmentation is essential to automatically extract features like organoid volume or structure. As shown in Figure 1a-b, the 3D U-Net reached an overall Dice score of 0.92±0.06 (mean ± SD) for organoid segmentation. Even though the model performs very accurately overall, we investigated challenging samples to identify the model’s weaknesses. The model performs poorest for Organoid 3 on day 36 (Dice score of 0.59). For this organoid, the disruption of one or more cystic structures resulted in a reduced overall volume (Supplementary Figure 1) and a split of the organoid into multiple pieces (Figure 1c). These pieces stick to the Eppendorf tube wall which causes that part of the organoid border blurs with the MRI background. This biological outlier is unique in our dataset and was therefore difficult to be learned by the model. The analysis of other samples shows that the model captured the organoids very well (Figure 1d-e).

**Figure 1.**
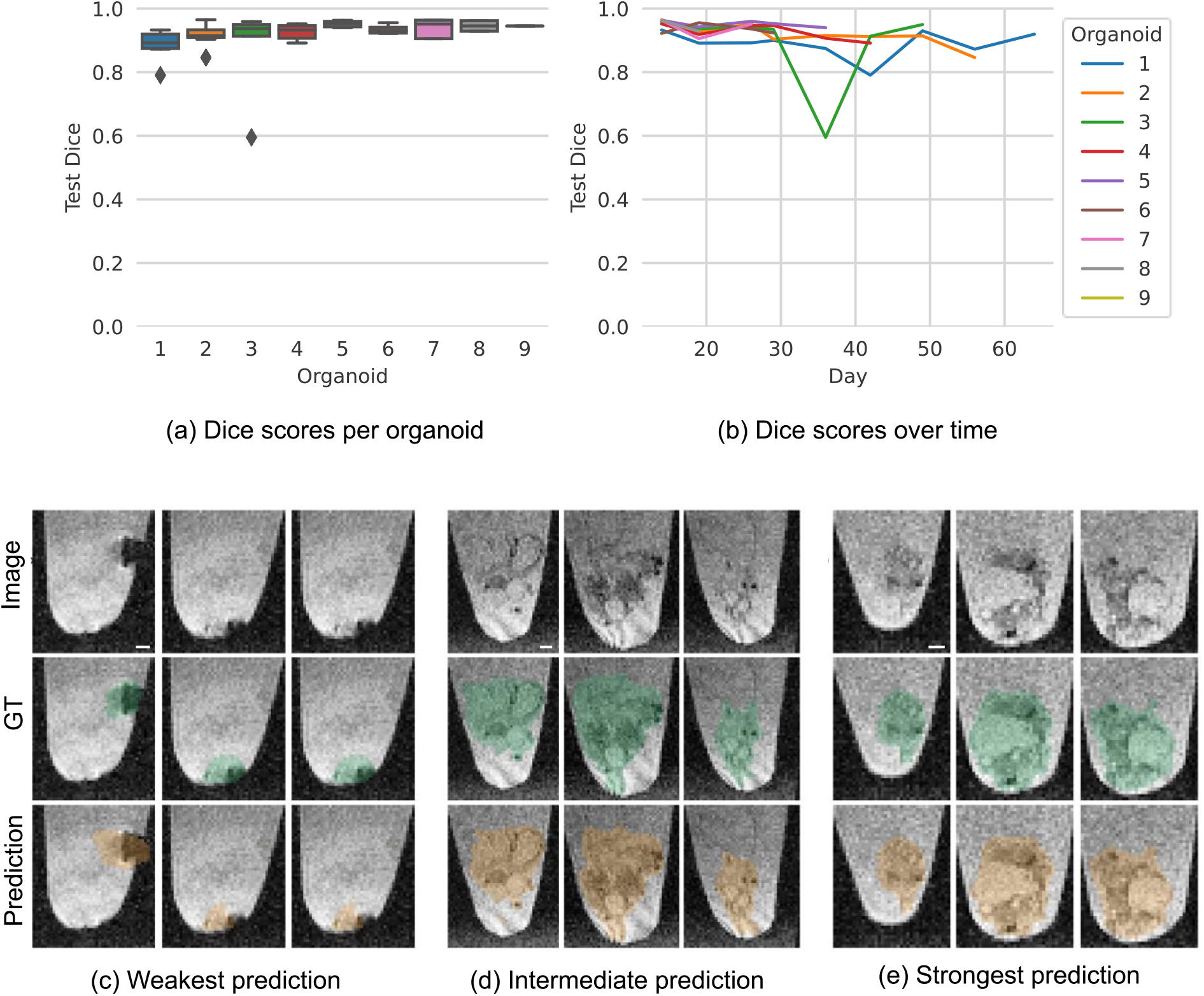
Organoid segmentation. (a) - (b) Model performance. (c) - (e) Selected sagittal planes. (c) Organoid 3 (day 36): Dice score of 0.59. (d) Organoid 2 (day 42): Dice score of 0.91. (e) Organoid 5 (day 26): Dice score of 0.95. Image: original image, GT: Image with ground truth organoid location (green), Prediction: image with predicted organoid location (orange). Selected sagittal planes (left to right): (c) 50, 47, 44 (d) 58, 50, 40 (e) 52, 40, 34. For better visibility, we cut the images to the Eppendorf tube boundaries. Scale bar: 400 µm.

### Global cysticity classification

Cyst formation is an undesired process during cerebral organoid cultivation [7]. Thus, accurately determining organoid cysticity can serve as a quality control tool. Separating low- and high-quality organoids using only their mean intensities resulted in a ROC AUC of only 0.65. However, our metric *compactness* achieved a ROC AUC of 0.98 (Figure 2).

**Figure 2.**
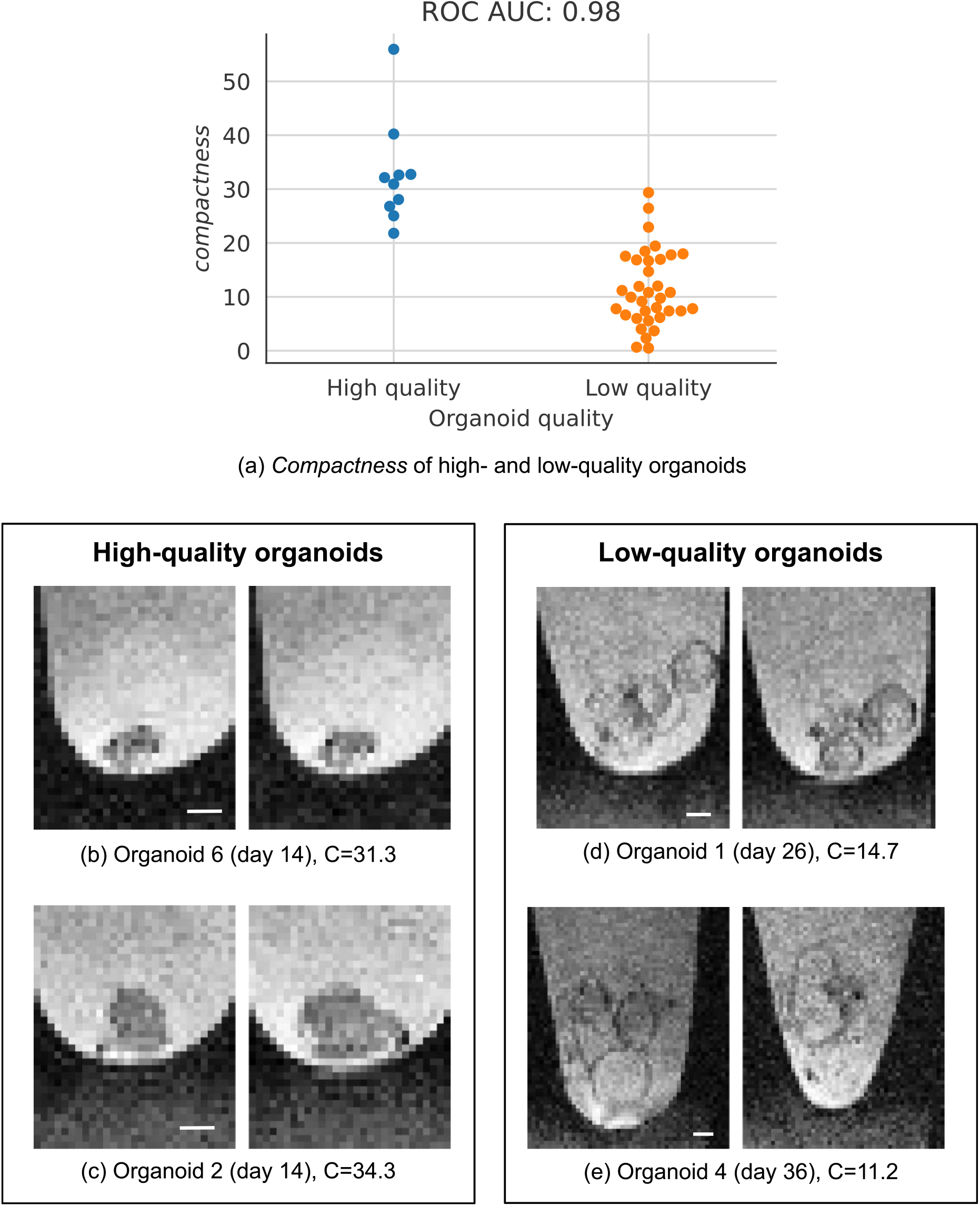
Global cysticity classification. (a) *Compactness* separates high- and low-quality organoids. (b) - (e) Selected sagittal planes from two low- and two high-quality organoids. C = *compactness*. Selected sagittal planes (left to right): (b) 59, 60 (c) 41, 45 (d) 58, 61 (e) 36, 53. For better visibility, we cut the images to the Eppendorf tube boundaries. Scale bar: 400 µm.

Using diffusion tensor imaging (DTI), we observed that low-quality organoids have a significantly higher average diffusion than high-quality organoids (Figure 3a). As can be seen in Figure 3b-c, cysts have an increased diffusion compared to compact tissue. Analysis of other parameter maps are included in Supplementary Table 1.

**Figure 3.**
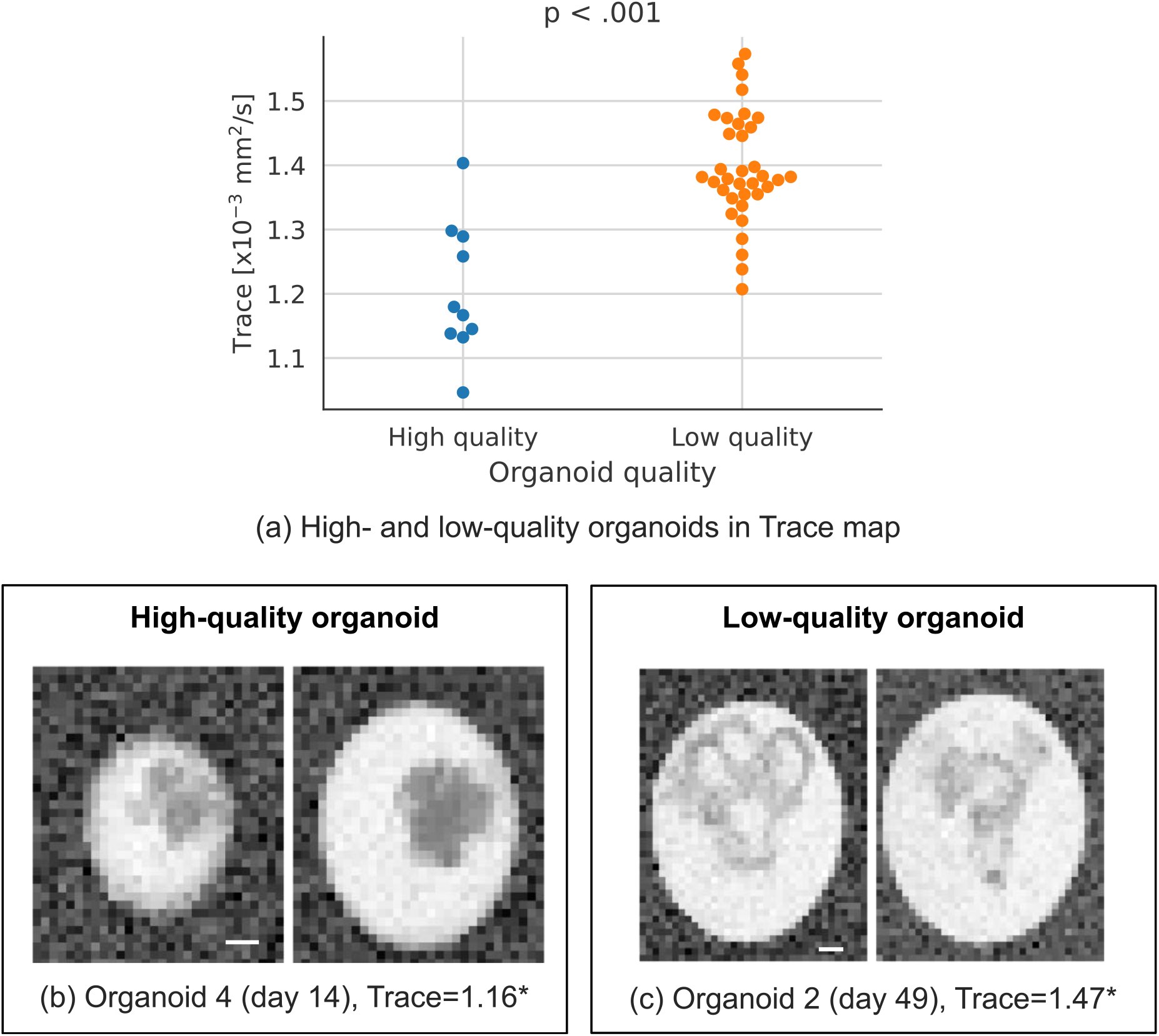
Diffusion tensor imaging (Trace map) shows different tissue characteristics of low- and high-quality organoids. (a) Trace of high- and low-quality organoids; p-value: two-sided t-test, adjusted with Holm-Šídák for multiple hypothesis testing. (b) - (c) Selected coronal planes from one low- and one high-quality organoid; *[x10^−3^ mm^2^/s]. Selected coronal planes (left to right): (b) 1, 2 (c) 5, 6. For better visibility, we cut the images to the Eppendorf tube boundaries. Scale bar: 400 µm.

### Local cyst segmentation

The good performance for global cysticity classification raises the question of whether cysts can be segmented locally – which would provide further insight into cyst distribution and location. For this task, the 3D U-Net achieved an overall Dice score of 0.63±0.15 (mean ± SD). As shown in Figure 4a-b, the Dice scores for individual samples showed a large variation with values ranging from 0.34 to 0.83. The analysis of weak and intermediate model predictions showed discrepancies between model predictions and ground truth especially for organoids with many small cysts (Figure 4c-d). The model performed especially well on images with large, clearly visible, and distinct cysts (Figure 4e).

**Figure 4.**
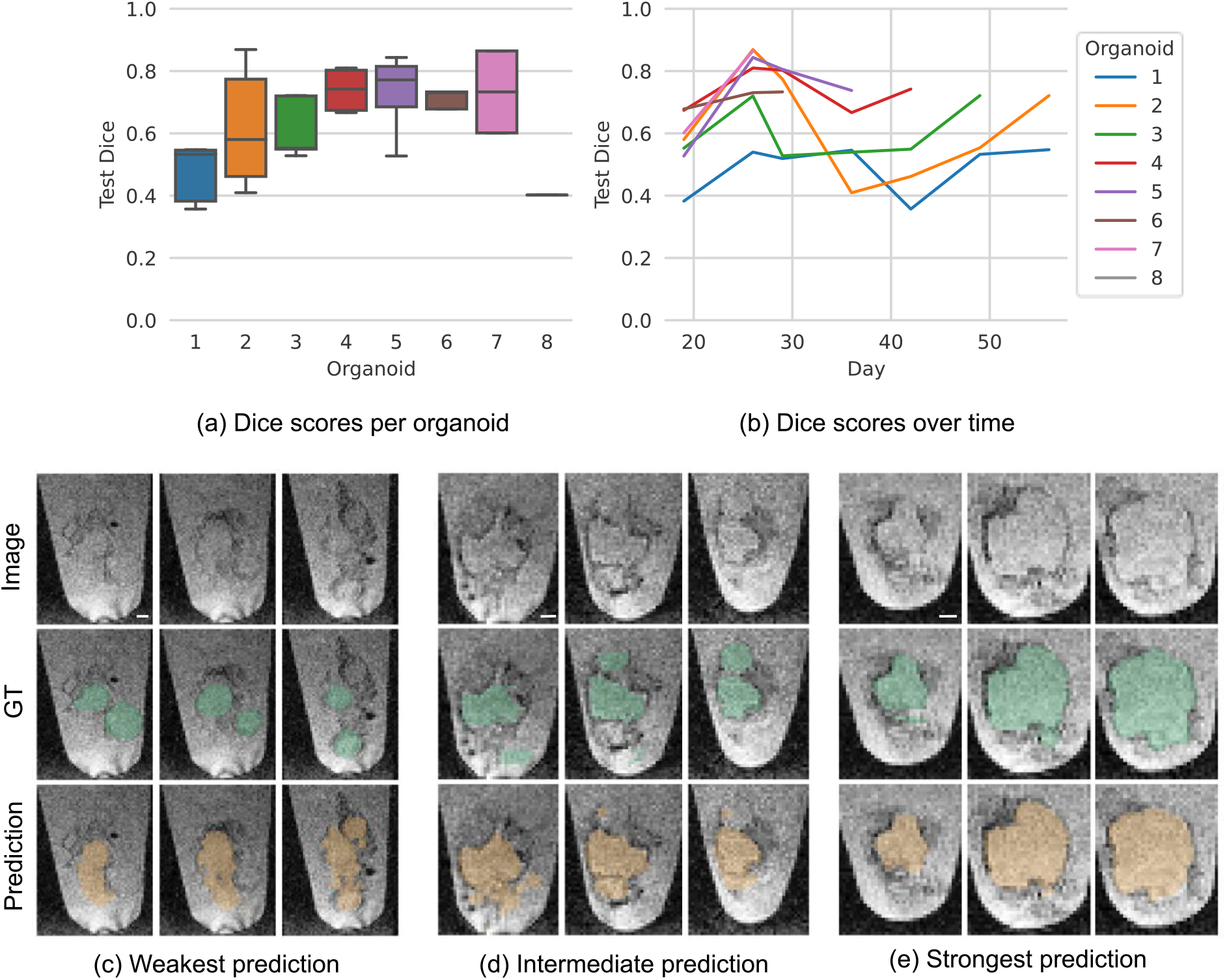
Local cyst segmentation. (a) - (b) Model performance. (c) - (e) Selected sagittal planes for three organoids. (c) Organoid 1 (day 42): Dice score of 0.34. (d) Organoid 4 (day 36): Dice score of 0.63. (e) Organoid 7 (day 26): Dice score of 0.83. Image: original image, GT: image with ground truth organoid location (green), Prediction: image with predicted organoid location (orange). For better visibility, we cut the images to the Eppendorf tube boundaries. Selected sagittal planes (left to right): (c) 60, 55, 51 (d) 52, 49, 42 (e) 63, 56, 49. Scale bar: 400 µm.

## 3. Discussion

In this study, we introduce high-field MRI for the non-invasive monitoring and analysis of cerebral organoids using a neural network-based approach. Since neither thresholding nor using a 2D U-Net resulted in convincing results for organoid segmentation (Supplementary Table 2), we used a 3D U-Net which achieved a mean Dice score of 0.92 for organoid segmentation. Comparable methods for MRI brain segmentation achieve Dice scores in the range of 0.72 and 0.93 [17-22]. Such a highly reliable automated analysis will represent a powerful tool to compare wild-type organoids with disease models associated with altered growth rate such as Zika-Virus disease [23] or microcephaly [24].

As the first step, reliable organoid segmentation paves the way for comprehensive quality monitoring including morphological and functional tissue parameters. The newly introduced metric *compactness*, inspired by the concept of signal-to-signal ratio [25, 26], assesses overall cysticity. It successfully separated high- and low-quality organoids at an outstanding ROC AUC of 0.98, closely matching the phenotypical appearance of previously reported high- and low-quality organoids [7, 8]. On a functional level, as expected, it was shown that low-quality organoids have a significantly higher diffusion than high-quality organoids most likely reflecting higher fluid content.

Successful global cysticity assessment led to the question of whether cysts can be segmented locally to differentiate solid compartments from fluid-filled cavities. The 3D U-Net trained for local cyst segmentation reached a mean Dice score of 0.63 which indicates a challenging segmentation task. Other challenging segmentation tasks such as ischemic stroke lesion segmentation achieve even lower Dice scores of 0.37 in MRI [27, 28] and 0.54 in CT [27, 29]. Especially for organoids having many small cysts, correct local cyst segmentation appears to be a major challenge due to technical resolution and contrast-to-noise ratio limits. In such cases, global cysticity classification may thus capture more easily the fluent transition from compact to cystic organoids.

Some limitations need to be taken into consideration. On the one hand, reliable organoid segmentation and global cysticity assessment could be achieved despite the relatively small dataset and heterogeneous organoid morphology. Thus, we do not expect a boost in performance here when extending the dataset. On the other hand, local cyst segmentation could probably benefit from a larger dataset. However, technical limitations of the image acquisition would most likely still impede segmentation performance in case of many small cysts due to uncertainty with respect to exact boundary detection for both human annotation and model prediction.

Overall, this work presents the first application of MRI for the non-invasive analysis of cerebral organoids. It was shown that cerebral organoids can be accurately monitored over time and for quality assessment using state-of-the-art tools for automated image analysis. These results highlight the potential of our pipeline for clinical application to larger-scale comparative organoid analysis.

## 4. Materials and Methods

The code to reproduce the results is publicly available on GitHub (https://github.com/deiluca/cerebral_organoid_quant_mri). All MRI images and annotations for organoid segmentation, global cysticity classification, and local cyst segmentation generated for this work are publicly available on Zenodo (https://zenodo.org/record/7805426, DOI: 10.5281/zenodo.7805426).

### Differentiation of cerebral organoids

Organoids were generated according to [10] with minor modifications. Wildtype iPSCs were singled and seeded at a density of 8×10^4^ cells/ml in a V-shaped 96 well plate in organoid formation medium (DMEM/F12, KnockOut Serum Replacement, NEAA, ß-mercaptoethanol) supplemented with 4ng/ml bFGF and Y-27632 (50µM) to induce embryoid body (EB) formation. The following day, the medium was exchanged to remove Y-27632 and lower the bFGF concentration to 2ng/ml. On day 5, neural induction was initiated by exchanging the medium to neural induction medium (DMEM/F12, N2 supplement, NEAA, glutamine, 1 µg/ml heparin) with a medium change on day 7. On day 9, EBs were embedded into Matrigel droplets and cultivated until day 13 in organoid differentiation medium (ODM) 1 (DMEM/F12:Neurobasal medium 50:50, NEAA, glutamine, penicillin/streptomycin, N2 supplement, B27 supplement w/o vitamin A, insulin, ß-mercaptoethanol). On day 13, organoids were excised from the droplets and transferred into a 12-well plate containing organoid differentiation medium 2 (DMEM/F12:Neurobasal medium 50:50, NEAA, glutamine, penicillin/streptomycin, N2 supplement, B27 supplement with vitamin A, insulin, ß-mercaptoethanol) and placed on a shaker in the incubator with medium exchange every 2-3 days. After imaging, the organoids were transferred back to the plate containing fresh medium and placed on the incubation shaker for further development.

### MRI

For MR measurements, organoids were transferred to 1.5 ml Eppendorf tubes containing standard ODM (T2-time of ∼64 ms in this experimental setting) and conveyed to the MRI using warming packs for temperature control. In total, nine organoids were scanned at varying time points over a period of 64 days, resulting in 45 individual samples. Three Eppendorf tubes were placed next to each other in a holder, thus allowing simultaneous imaging of three organoids (Supplementary Figure 2). Nine control organoids not undergoing MRI served as handling control. Before and after imaging, the medium was analyzed in both groups using a blood gas analyzer which showed that MRI had no specific negative effect on organoid health (Supplementary Table 3).

MRI was performed at room temperature using a high-field 9.4 Tesla horizontal bore small animal experimental NMR scanner (BioSpec 94/20 USR, Bruker BioSpin GmbH, Ettlingen, Germany) equipped with a four-channel phased-array surface receiver coil. The MR protocol included the following sequences:

1. High-resolution T2*-weighted gradient echo sequence: 3D sequence, echo time (TE): 18 ms, repetition time (TR): 50 ms, 80 µm isotropic resolution, acquisition matrix: 400 × 188 × 100, flip angle: 12°, number of averages: 1, duration: 15 min 40 s. This sequence was chosen to allow for accurate isotropic imaging and to account for potential susceptibility effects caused by e.g. neuromelanin [11], cellular debris or calcifications.
2. DTI-spin echo sequence: 2D sequence, TE: 18.1 ms, TR: 1200 ms, 100 µm in-plane resolution, acquisition matrix: 120 × 50, slice thickness: 1.5 mm, number of diffusion gradient directions: 18 + 5 A0 images, b-values: 0/650 s/mm^2^, gradient duration: 2.5 ms, gradient separation: 15.5 ms, flip angle: 130°, number of averages: 1, duration: 23 min 05 s. This sequence was included to account for organoid inner structure including nerve fiber growth [12].

### Organoid segmentation

Organoid segmentation was performed to assign each image voxel to one of two categories: organoid or non-organoid. For this task, we used min-max normalized images from the T2*-w sequence. Since simpler methods like Multi-Otsu’s threshold [13] and a 2D U-Net [14] did not deliver convincing results (Supplementary Table 2), we used a 3D U-Net [15] for efficient (Supplementary Table 4) organoid segmentation. We trained the model with Adam (learning rate 1×10^−3^, weight decay 1×10^−7^) for 2,000 iterations with batch size 1 and a combination of binary cross entropy and Dice loss.

For model evaluation, we used the Dice score, which is commonly used to quantify the performance of image segmentation methods. It is defined as two times the area of the intersection divided by the total number of voxels in the ground truth and predicted segmentation (Eq. 1). A perfect segmentation corresponds to a Dice score of 1.

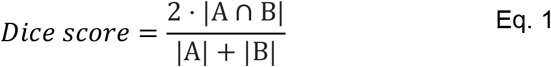

To get an unbiased estimate of the model performance, we used organoid-wise Leave-One-Out Cross-Validation (LOOCV). For each of the nine LOOCV splits, we used a random 80% training, 20% validation split for model selection. The Dice score in the Results section refers to the model performance on the LOOCV test set.

### Global cysticity classification

Global cysticity classification aims at determining the overall organoid cysticity: cystic (low-quality) or non-cystic (high-quality). To provide a reference ground truth based on the T2*-w sequence, an organoid was categorized as low-quality if a cystic structure was detected within the organoid, consistent with findings on brightfield imaging (Supplementary Figure 3) as previously reported [7, 8]. Otherwise, it was categorized as high-quality.

For automatic classification, we constructed the simple metric *compactness* which serves as an environment-based estimator of organoid cysticity (Eq. 2). It is based on the idea that cysts are filled with similar fluid like the medium under the assumption of relative B1-homogeneity in a stereotyped region close to the surface coil. Therefore, the more similar the organoid intensities are to the medium intensities, the more cystic the organoid is.

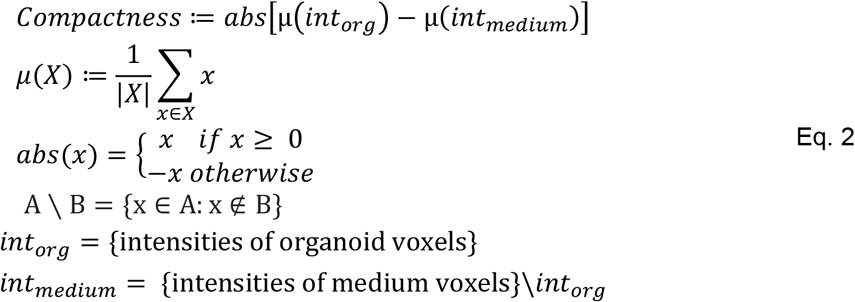

While *int*_*org*_ was derived from the ground truth organoid segmentations, *int*_*medium*_ was determined by applying Otsu’s threshold [16] 2D-wise along all organoid-containing coronal planes (Supplementary Figure 4). The first and last organoid-containing coronal planes were discarded to filter artifacts caused by noisy medium intensities.

For the evaluation of *compactness*, we used the area under the Receiver Operator Characteristic curve (ROC AUC). ROC AUC is a common metric for the evaluation of binary classification problems; a perfect classifier achieves a ROC AUC of 1.

To further probe tissue characteristics of low- and high-quality organoids, parameter maps (Trace; FA; 1^st^, 2^nd^, and 3^rd^ Eigenvalues) were extracted from the DTI sequence using the built-in analysis tool (Paravision 6.0, Bruker BioSpin GmbH, Ettlingen, Germany). We used a two-sided T-test to test for significantly different average diffusion and used Holm-Šídák to adjust for multiple testing.

### Local cyst segmentation

Local cyst segmentation aims at localizing cysts. For this task, we used the T2*-w sequence and manually annotated cysts. Due to the low-resolution images, especially smaller cysts are difficult to annotate. Therefore, we excluded organoids with less than 1,000 voxels (0.51 mm^3^) in cysts and included 34 samples in total. For segmentation, we trained and evaluated a 3D U-Net [15] as for organoid segmentation but with 5,000 training iterations.

## Supplementary Figures

**Supplementary Figure 1.**
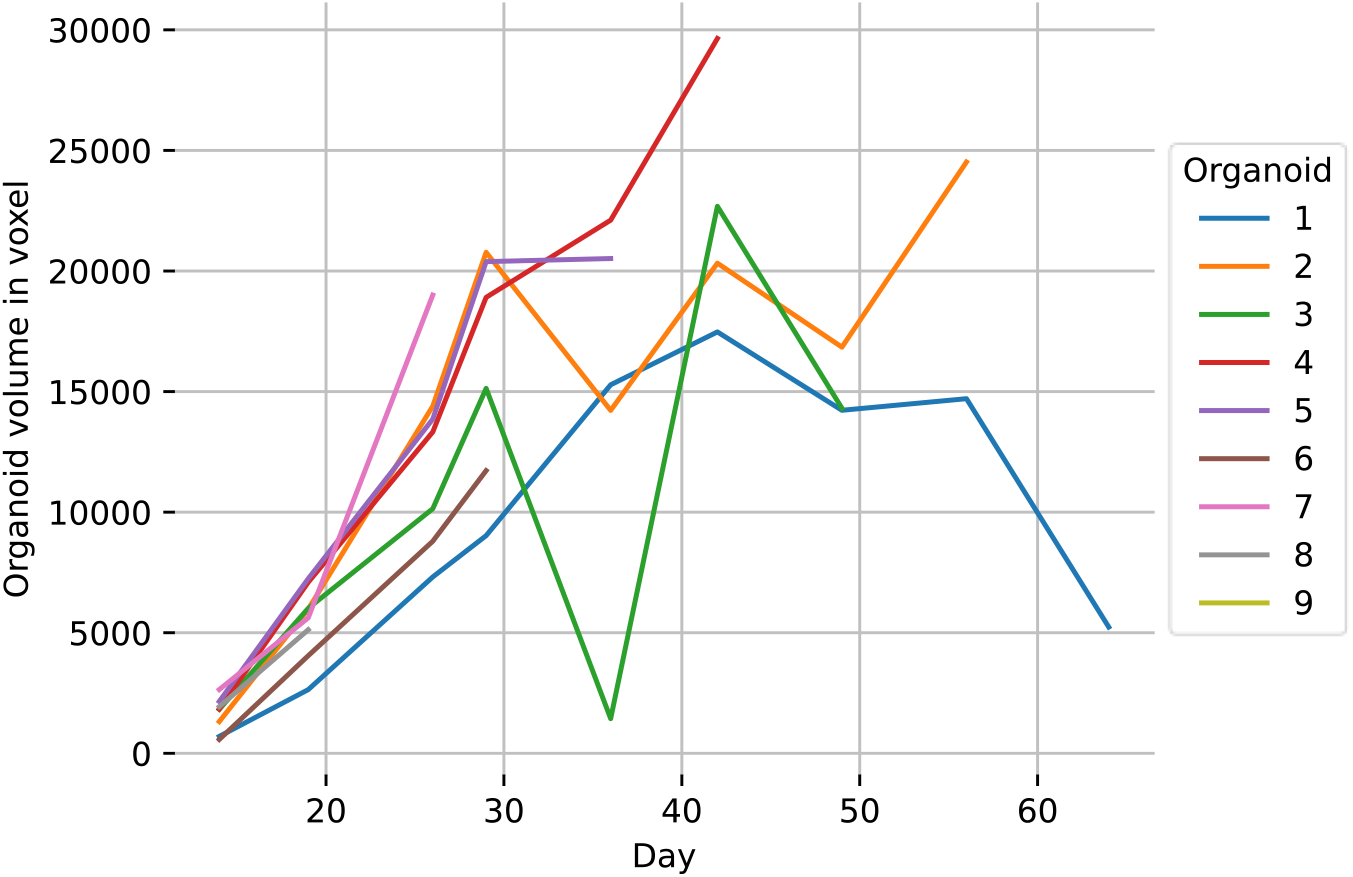
Organoid growth over time. The organoid volume in voxels (1 voxel = 5.12 × 10^−4^ mm^3^) is based on the ground truth organoid annotation in the T2*-w sequence. Organoid 3 (day 36) has a sudden drop in volume which is due to the disruption of one or more cystic structures. Exemplary planes of this organoid are shown and discussed in the main text.

**Supplementary Figure 2.**
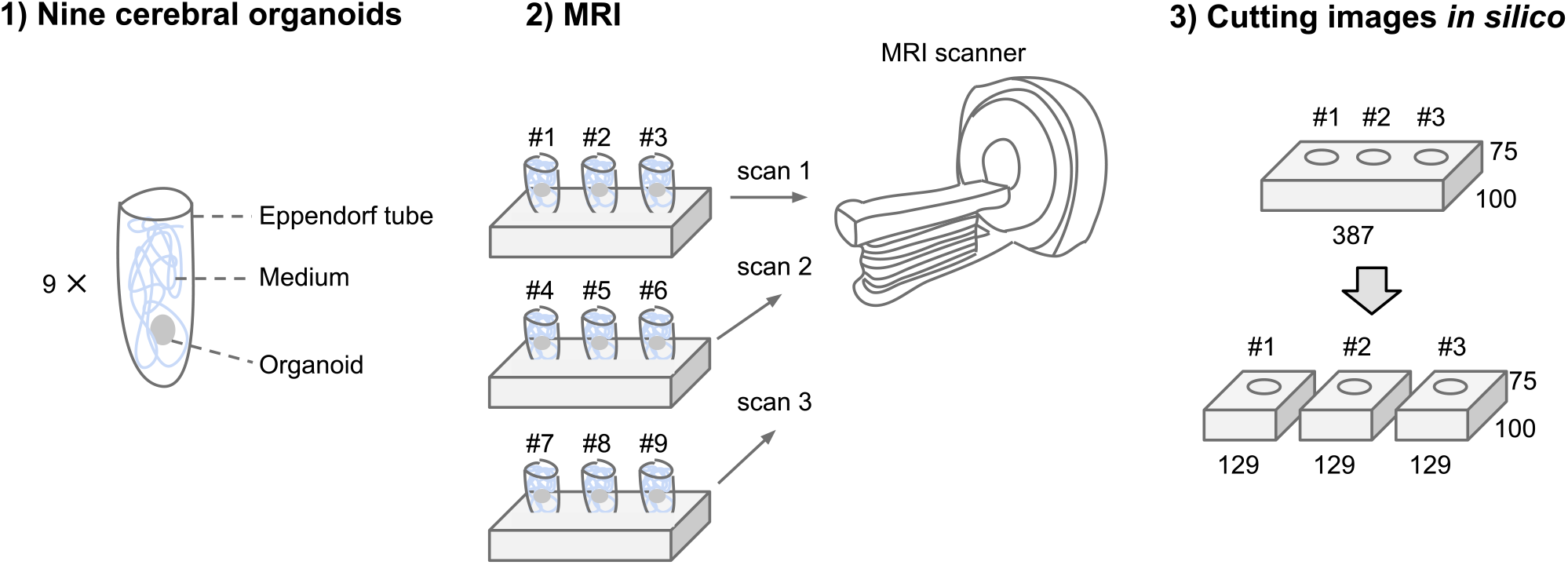
Experimental setup and data acquisition. For MRI, three Eppendorf tubes were placed next to each other in a holder. Subsequently, the images were cut *in silico* to derive one image per organoid. Image dimensions shown in 3) are according to the T2*-w sequence.

**Supplementary Figure 3.**
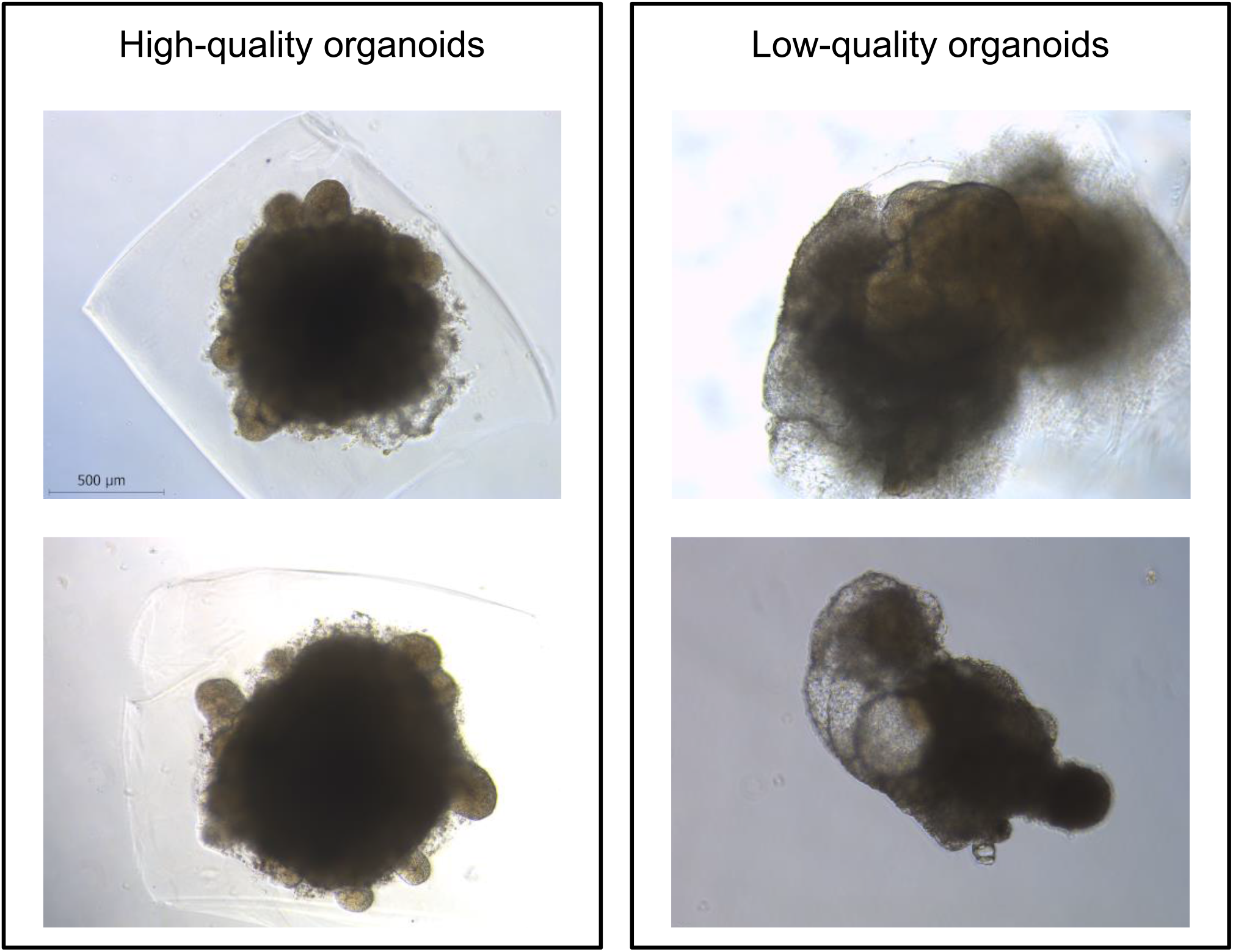
Microscopy images of two high-quality (non-cystic) and two low-quality (cystic) organoids on day 14 of organoid differentiation. The low-quality organoids show fluid-filled cavities (or “cysts”) and therefore resemble the same phenotype as reported in [7, 8].

**Supplementary Figure 4.**
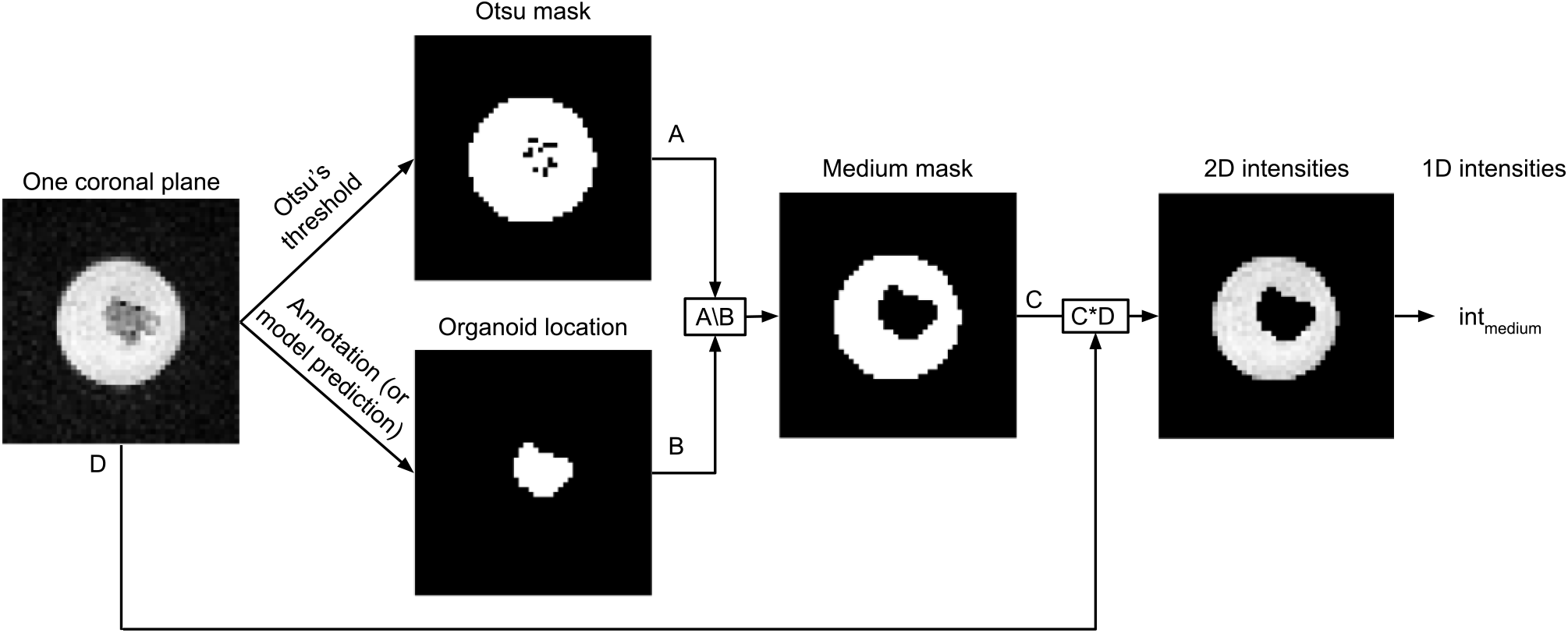
Concept of medium intensity determination for global cysticity classification. Otsu’s mask, organoid location and medium mask are binary masks. The white pixels of the medium mask belong to the medium. This example is based on Organoid 1 (day 14), coronal plane 60. To determine the medium intensities for one organoid, this procedure is applied to all organoid-containing coronal planes. For better visibility in this figure, we cut the coronal plane to the Eppendorf tube boundaries.

## Supplementary Tables

**Supplementary Table 1.** ROC AUCs and adjusted p-values for separation of low- and high-quality organoids for selected DTI parameter maps.

| DTI parameter map | ROC AUC | P-value |
| --- | --- | --- |
| Trace | 0.91 | $1.1 \times 10^{-5}$ |
| 3rd Eigenvalue | 0.86 | $6.2 \times 10^{-4}$ |
| 2nd Eigenvalue | 0.91 | $1.2 \times 10^{-5}$ |
| 1st Eigenvalue | 0.93 | $2.1 \times 10^{-5}$ |
| Fractional Anisotropy (FA) | 0.63 | $9.9 \times 10^{-1}$ |

**Supplementary Table 2.**
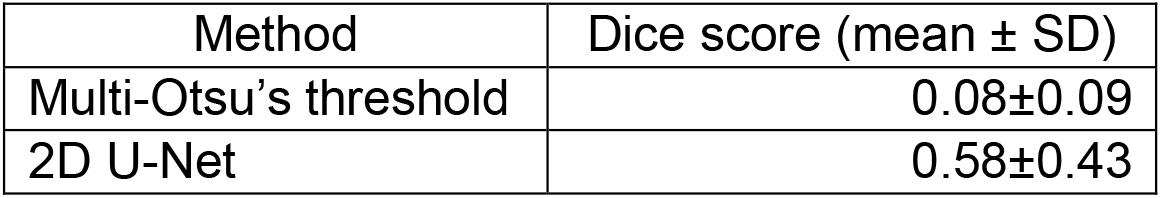
Organoid segmentation performance of Multi-Otsu’s threshold [13] and 2D U-Net [14] in the T2*-w sequence. Multi-Otsu’s threshold was applied in 3D for the three classes MRI background, Eppendorf tube, and organoid using the Python package scikit-image. For the 2D U-Net, the images were extracted along the coronal axis. For 2D U-Net training and evaluation, the implementation from https://github.com/milesial/Pytorch-UNet was utilized. 2D U-Net: binary semantic segmentation; 200 epochs; batch size 1; learning rate 0.00001; loss: binary cross entropy + Dice loss (weighted 1:10), weight decay: 0.001; augmentation: random rotation (probability 0.75).

**Supplementary Table 3.** Blood gas analysis shows no specific negative effect of MRI on organoids. Median differences of all pre- and post-MRI measurements for medium control w/o organoid (‘Medium’), MRI organoids (Org_MRI_), and control organoids (Org_control_). Cells are colored according to measurement increase or decrease.

| Measurement | Medium | Org<sub>MRI</sub> | Org<sub>control</sub> |
| --- | --- | --- | --- |
| pH Medium | 0.02 | -0.29 | -0.31 |
| pCO<sub>2</sub> [mmHg] | -0.65 | 14.30 | 15.70 |
| pO<sub>2</sub> [mmHg] | -4.10 | 2.30 | -6.40 |
| HCO<sub>3</sub><sup>-</sup> act [mmol/l] | -0.35 | -2.60 | -2.50 |
| HCO<sub>3</sub><sup>-</sup> std [mmol/l] | 0.50 | -7.20 | -7.50 |
| Glucose [mg/dl] | -7.00 | -22.00 | -24.00 |
| Na<sup>+</sup> [mmol/l] | 0.20 | 1.10 | 1.70 |
| K<sup>+</sup> [mmol/l] | 0.00 | 0.02 | 0.04 |
| Ca<sup>2+</sup> [mmol/l] | -0.01 | 0.00 | -0.01 |
| Cl<sup>-</sup> [mmol/l] | 0.00 | 1.00 | 1.00 |

**Supplementary Table 4.** Efficient 3D U-Net training and inference for organoid segmentation. For application to larger-scale experiments, it is important that the model training and especially inference time are in a practical range. The 3D U-Net requires less than an hour for training on MRI organoid segmentation using the T2*-w sequence. Inferring the model predictions is in the range of two seconds per sample. The times were measured using one NVIDIA GeForce RTX 3090 (24 GB) graphics card.

| Model | Training time (s) |  | Inference time (s) |  |
| --- | --- | --- | --- | --- |
|  | Per iteration | Total | Per sample | Total |
| 3D U-Net | 1.11 | 2,220 | 1.97 | 88.6 |

## Notes

https://zenodo.org/deposit/7805426

https://github.com/deiluca/cerebral_organoid_quant_mri

### Competing Interest Statement

The authors have declared no competing interest.

### Summary of Updates

Now Results before Materials and Methods; Previous Figure 1 and 2 now in Supplementary Figures; Radiology-specific abbreviations introduced.

